# Never Mind the Repeat: How Speech Expectations Reduce Tracking at the Cocktail Party

**DOI:** 10.1101/2025.03.21.644185

**Authors:** Thaiz Sánchez-Costa, Alejandra Carboni, Francisco Cervantes Constantino

## Abstract

When the brain focuses on a conversation in a noisy environment, it exploits past experience to prioritize relevant elements from the auditory scene. This prompts the question of what changes occur in the selective neural processing of speech mixtures as listeners garner prior experience about single speech objects. In three different priming experiments, we quantified cortical selection of temporal landmarks from continuous speech, applying the temporal response function (TRF) method to single-trial electroencephalography (EEG) recordings. The designs specifically addressed how attention interacts with exact (Experiment 1), voice (Experiment 2a), or message (Experiment 2b) content priming of the target or background speakers in cortical responses to speech. Our results demonstrate that, during multispeaker listening, attentional gains typical of cortical responses under speech selection are met with attenuations as a consequence of prior experience. The changes were observed at the P2 processing stage (220-320 ms) of speech envelope onset processing and were specific to responses to primed speech targets (Experiment 1). Suppressions at stages earlier than the P2, or under partial priming conditions (Experiments 2a and 2b), were not observed. An exploratory analysis suggests the observed P2 reduction predicts listeners’ ability to report target words, consistent with this component encoding in part temporal prediction error about onset edge cues exclusive to target speech. Our results show that at this late and definitive stage of selective attention, the auditory system may test the evidence for its own predictive model of the noise-invariant speech stream. Precise inference of its temporal structure is bound to tag all checkpoints where auditory evidence can be most reliably connected into higher-order representations of continuous speech.

## Introduction

Social living requires communicating and understanding. Soundscapes carry background noise, yet listeners often retain the ability to choose and follow target speakers. Selective attention operates on this “cocktail party” scenario (<u>Cherry, 1953</u>), where the peripheral representation of mixed sound is incrementally shaped for the brain to single out and parse its target ( <u>Har-shai Yahav and Zion Golumbic, 2021</u>; <u>Wood and Cowan, 1995</u>). Analyzing the target from the rest of the scene in a behaviourally meaningful way requires that parallel neural processes feedback upon one another, and part of this balance relies on the observer’s experience and prior expectations (<u>Shinn-Cunningham, 2008</u>; <u>Shinn-Cunningham et al., 2017</u>). For example, if we have to select and understand somebody amid other talkers, knowing their particular voice well will likely help us (<u>Domingo et al., 2020</u>). Voice familiarity may also prove advantageous when we must ignore the same person’s speech ( <u>Newman and Evers, 2007</u>; <u>Johnsrude et al., 2013</u>). F oreknowledge about speech content is another type of behaviourally relevant experience in the cocktail party ( <u>Dekerle et al., 2014</u>; <u>Park et al., 2023; Bhandari et al., 2021</u>). Indeed, exact or full prior knowledge about a target represents the ground truth condition to evaluate experience biases on the balance of neural processing, which favors segregation amid maskers (<u>Wang et al., 2021</u>). Yet the precise stage at which such changes first occur in the brain remains unresolved.

The representations of a speech stream, which emerge across cortical networks during listening may be investigated using temporal response function (TRF) methods in combination with high temporal precision recordings (<u>Bednar and Lalor, 2020</u>; <u>Broderick et al., 2019</u>; <u>Di Liberto et al., 2015</u>; <u>Har-shai Yahav and Zion Golumbic, 2021</u>; <u>Wöstmann et al., 2019</u>; <u>Zion Golumbic et al., 2013</u>). The TRF technique delineates how much and when the brain is likely to engage with specific acoustical characteristics that are present in the masked speech signal (<u>Ding and Simon, 2012</u>; <u>Haykin and Chen, 2005</u>; <u>O’Sullivan et al., 2015</u>; <u>Power et al., 2012</u>; <u>Fiedler et al., 2019</u>; <u>Ding and Simon, 2012</u>; <u>Kidd et al., 2016</u>). For instance, it is used to describe the ability of auditory responses to align with temporal regularities found in the speech envelope, a phenomenon known as ‘phase coding’ or ‘speech tracking’ (<u>Obleser and Kayser, 2019</u>; <u>Zion Golumbic et al., 2013</u>). This stimulus-locked response is often organized into a triphasic neural entrainment pattern whose individual stages typically correspond to the P1, N1, and P2 temporal components of the auditory event-related potential and are likewise named (<u>Aiken and Picton, 2008</u>; <u>Fiedler et al., 2019</u>; <u>Martin et al., 2008</u>; <u>Steinschneider et al., 2011</u>). For the cocktail party, attentional gains appear reliably in auditory responses to the target speech around 100 ms, corresponding to the N1 component of the TRF (<u>Ding and Simon, 2012</u>).

Does prior experience synergistically boost or antagonistically decrease selective enhancement at this point? In sensory systems, unpredicted stimuli may carry surprise signals for which additional processing is often involved. Auditory surprise signals may be differentially coded by attentional selection, as evidenced by the boosting of cortical mismatch responses to frequency deviant tones by temporal attention around 200 ms post onset (Auksztulewicz and Friston, 2016). In the cocktail party, it is not yet clear whether attention effects and expectations arising from prior experience analogously interact at the N1 or other relevant stage of speech processing. Unlike mismatch sequence paradigms, in the cocktail party the object streams meet different attention and expectation conditions to be accounted for simultaneously. Moreover, the focus of mismatch designs is the novelty gain response (which attention may boost), while in the cocktail party the focus is the attentional gain response (which repeated experience could conversely reduce). Determining how both interact and the stage of speech processing at which they may do so matters in the context of perceptual inference. In these problems, inference is driven by observer expectations generating internal predictions to be compared against input signals from bottom-up sensory input (<u>Clark, 2013</u>; Ten Oeve & Martin, 2024). The stage of the first interaction and the precise aspects of speech that are engaged by prior experience may hence single out the learning processes by which the auditory brain shapes objects in the cocktail party.

In a magnetoencephalography (MEG) study, <u>Wang et al.</u> (<u>2019</u>) directly addressed whether exact prior knowledge of a speech stream modulates selective locking responses in the auditory cortex during cocktail party masking conditions. Under target priming, an attentional bias in favor of target-locked responses was observed and explained by weakened representations of masker speech, boosting in effect attentional gains relative to unprimed conditions. Because maskers were not primed, the findings appeared at odds with activity reductions that result from more efficient encoding of redundant information under expected stimuli (Grill-Spector et al., 2006; <u>Summerfield et al., 2008</u>). Expectation-related reductions featuring in speech processing include evidence that visual predictions decrease the amplitude of listeners’ N1 and P2 component auditory responses to corresponding speech (<u>Pinto et al., 2019</u>), for instance. Such research indicates that response reductions under priming may not even require exact sensory repetitions, relying instead on input that is redundant, accurate, or consistent with prior expectations of the observer. Altogether, these aspects raise the issue of whether priming information from a speech object specifically alters its representation during redundant recoding in the cocktail party. At the same time, they underscore the currently unknown issue of whether prior expectations similarly adjust the processing of speech objects that are to be ignored as distractors. This latter scenario may serve to address the continuum between active inhibitory processes arising from the intentions of the observer versus those that stem from the extraction of regularities over time through learning (<u>Wöstmann et al., 2022</u>).

In the present study, the interaction between selective attention and prior experience in speech processing was systematically addressed in the context of a speech comprehension task that represents typical challenges facing the ‘cocktail party’ problem. Using single-trial EEG and TRF tools, we investigated their interaction at each of the relevant P1, N1, and P2 stages of speech processing. In particular, we focused on the impact of auditory priming on the selective tracking of a critical cue in the segmentation and parsing of the temporal structure of speech, namely its temporal envelope edges. These auditory features have been demonstrated to be a predominant basis for stream segregation (Brodbeck et al., 2018). Our aim was to understand how they are tracked in parallel along with the observer’s unfolding temporal predictions about speech. One candidate mechanism is through online comparisons between edge signals and observer expectations about them, where exact correspondence minimizes uncertainty about temporal aspects of a target object during selection.

## General Methods

### Ethics statement

The Ethics in Research Committee at the Faculty of Psychology, Universidad de la República Uruguay, approved the study. All participants gave their written consent in accordance with the guidelines of the Declaration of Helsinki (World Medical Association, 2009). As a token of appreciation for their time, the participants received a chocolate bar at the conclusion of the experimental session.

### Participants

Seventy-four subjects participated in one of three different experiments conducted to probe the interaction between attention and prior experience, via priming of either the exact content of a speech signal, its voice, or message, respectively (48 female; mean age 25.6 ± 4.9 SD; 11 left-handed; see information detailed at specific Methods sections, below). Subjects were native Rioplatense Spanish speakers who reported no hearing impairment or neurological disorder and normal or corrected vision. The participants completed the <u>Edinburgh Handedness Inventory</u>.

### Setup

Participants were seated 50 cm in front of a 40 cm size monitor (E. Systems, Inc., CA). For audio presentation, we used a Sound Blaster Z sound card (Creative Labs, Singapore) in combination with a Scarlett 414 sound interface (Focusrite Plc, UK) and high-quality Sennheiser HD 25 headphones (Sennheiser, Germany) to present sounds diotically. Neural recordings were made with BioSemi ActiveTwo 64-channel system with an Active two ERGO Opticlink (BioSemi, The Netherlands) to connect synchronous auditory signals to the ActiveTwo output system. The participants adjusted the presentation of the sound volume to a comfortable listening level before starting the computerized tasks.

### Stimuli

#### Single speaker stimuli

Auditory stimuli were obtained from two databases specifically constructed for this study (see Stimuli <u>Database 1, 2</u>, below) which involve hundreds of different single speakers. These database audios corresponded each to a brief (∼8-9 s) single speaker narration that, for the purposes of the study, were later combined into target-background two-speaker mixtures (see Cocktail party stimuli, below). Database audios consisted of short phrases narrated by a single speaker, and with annotated sentence keywords and topics. Each audio was additionally classified according to the apparent age and sex group of the speaker. MATLAB® software (MathWorks, Natick, United States) was used to convert audio recordings into diotic format by averaging, at a sampling rate of 44100 Hz. The low-frequency amplitude envelope of speech signals was obtained using Hilbert’s method, which was used to remove long silences; repeated filler words were similarly manually removed. To prevent the audible perception of clicks, 5 ms long cosine ramps were applied at the beginning or end of the stimulus and any excisions. All individual stimuli were normalized.

#### Cocktail party stimuli

A unique set of cocktail party (CP) stimuli was created per study participant. Each CP stimulus was generated from a selection of two solo speech stimuli from the same database, by adding the two streams using a 750 ms inter-stream onset asynchrony. This design was used for target instruction purposes, to ensure that listeners know which single speech stimulus to attend to (i.e., starting or interjecting speaker). The end time of every CP stimulus equalled the resulting mixture overlap end time. The root mean squared (RMS) intensity was equalized between the overlapping epochs of both streams, and the amplitude of the entire cocktail stimulus was normalized.

### Experimental task

We investigate the interaction between prior experience and attention, specifically how previous listening experiences may affect listeners’ neural tracking on a target speaker in a multi-talker scenario. For this, each trial followed a two-stage design and participants were instructed to listen to every auditory stimulus presentation with their eyes closed while minimizing motion. In the first part, ‘pre-listening’, participants were presented with a single-speaker speech stimulus. In the second part, ‘CP listening’, participants were presented with a two-speaker mixture and tasked to selectively attend to either the first (leading) or second (interjecting) speech stream in the mix. Leading or interjecting targets were visually cued immediately beforehand, and instructed with equal probability. At the end of CP listening, the trial continued with a questionnaire designed to evaluate participants’ comprehension of the target speaker. The task was to select a minimum of three options that apply to the target speech, based on a nine-option multiple choice format. Subsequent to participant’s response validation, the score feedback of the trial was provided to participants, along with their total accumulated score for the experiment (Figure 1).

**Figure 1.**
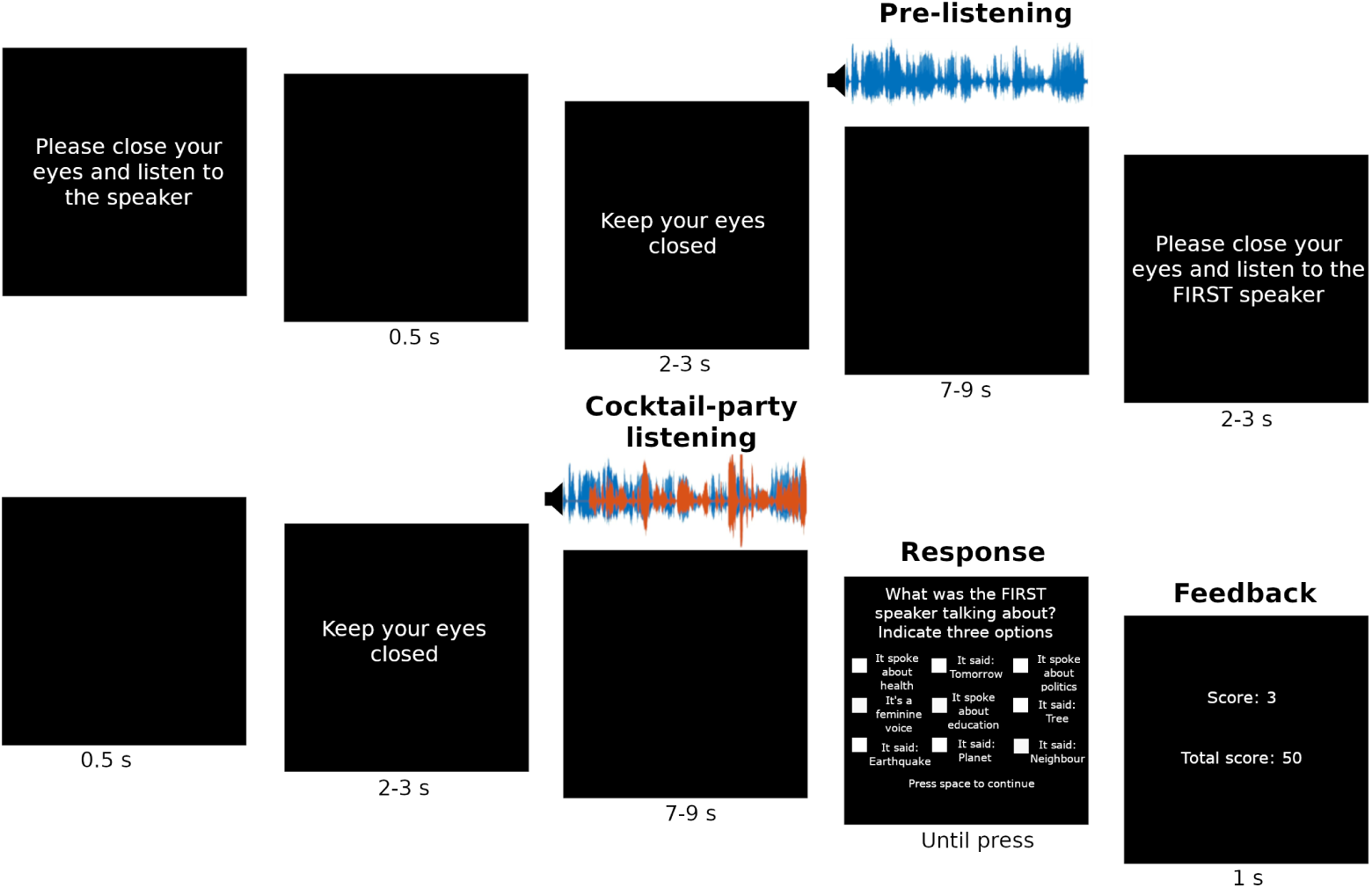
Experimental design. Each trial consisted of two parts, in the first, participants listened to a single speaker speech stimulus (pre-listening). In the second part, CP listening, participants were given instructions to attend to one speaker from a cocktail-party stimulus, i.e., a speech mixture of two different speakers. At the end of the presentation, participants were then required to select correct statements relating to the attended speech. The example shown illustrates the condition where the pre-listening stage provides exact prior knowledge about the target element of the CP mix (‘attended known’).

### Experimental conditions

Within each experiment, we defined three conditions to provide listeners with different degrees of knowledge about individual streams present at the CP listening stage, namely, attended known (AK), unattended known (UK), and no prior knowledge (NP). In AK[UK] trials, the pre-listening presentation matched speech information contained in the target[masker] stream of the CP listening mixture. In NP trials, the pre-listening presentation had no connection to either the target or masker stream of the CP listening mixture, serving as a control condition and reference across all experiments. Before the start of the CP-listening stage, listeners were indicated which stream to target based on the leading or interjecting order (“first/second”) of the asynchronous onset mixture (Figure 1, see Cocktail party stimuli). Every pre– and CP-listening stimulus was presented only once. Prior to the main experimental session, participants performed a three-trial practice session using fixed non-database stimuli that were specifically generated for this purpose. In all experiments, all speakers systematically varied from trial to trial, with random and unique stimuli combinations created per participant without stimulus replacement. All three conditions were equally represented and randomly ordered during the experimental sessions.

### Task questionnaire and scoring system

Participant behavior was assessed via a multiple-choice questionnaire that was presented after the CP-listening stage of each trial (Figure 1). Nine options were generated per trial, based on information specific to the target or masker stream stimulus and collected from their corresponding databases. One of the options was related to vocal characteristics, probing either the target speaker voice’s gender (male or female) or age (young or advanced age) attributes, with equal probability. Four of the additional options involved keywords that were present in the sentences, and they were randomly drawn from a list of all keywords from the target and masker stream. The remaining four options listed possible topics that the target stream speech may be categorized into, and were presented based on a random selection of our databases’ topic sets (see Stimulus Database 1, 2 for full details). In the case of topical information, it was possible that none of the presented categories corresponded to either stream. The placement of the nine different options was randomized for every trial.

For scoring, selecting an option that correctly related to the target speaker’s voice (i.e., sex or age) earned one point. Selecting an option that related to the masker speaker’s voice, and not the target’s, resulted in a point deduction. Similarly, each correctly chosen keyword from the target sentence awarded one point, and each selected keyword related to the masker sentence deducted one point. Each correct topic selection earned one point, but topic selections unrelated to the target sentence did not result in any point deductions. Participants received feedback on the trial score and on their accumulated score (see Figure 1).

### Data acquisition and preprocessing

Stimulus presentations were made with PsychoPy3 software (Peirce, 2007). Electroencephalography (EEG) data were recorded for all subjects using a BioSemi ActiveTwo 64 scalp channels system (BioSemi, The Netherlands) with 10/20 layout, at a digitization rate of 2048 Hz with CMS/DRL (ground) and the tip of the nose as reference electrodes. EEG subject data resampled at 1024 Hz were common average-referenced to the 64 scalp channels, after which DC offset was removed. A 5th order cascaded integrator-comb low-pass filter with –3 dB at 410 Hz was applied online, after which signals were decimated to 1024 Hz. The online high-pass response was fully DC coupled. Electrooculographic data were recorded, supra– and infra-orbitally as well as from the left versus right orbital rim. Complete experimental sessions lasted ∼ 2.5 h.

EEG data analyses were implemented in MATLAB 2018b. EEG signals were bandpass-filtered between 1 and 28 Hz with a 10-order elliptic IIR filter of 60 dB attenuation and 0.5 dB passband ripple, with group delay correction. Single channel data were subject to an automated blind rejection procedure based on a variance-based criterion (<u>Junghöfer et al., 2000</u>) using a confidence coefficient of *λ_P_*= 4. The procedure was performed separately for the external reference channels, with a more restrictive confidence coefficient *λ_P_*= 2. Sensor and reference data sets were separately re-referenced based on the channel sets median values, after which the channels rejection procedure was again conducted. FastICA (<u>Hyvarinen, 1999</u>) was applied and two independent components were automatically selected for their maximal proportion of broadband power, and projected out of the raw sensor data. A time-shifted principal component analysis (<u>de Cheveigné and Simon, 2008</u>) was applied to discard environmental signals recorded on the oculogram reference sensors (time shifts: ±4 ms). A sensor noise suppression algorithm (<u>de Cheveigné and Simon, 2008</u>) was applied to attenuate artifact components specific to any single channel, using 63 neighbors. Finally, the blind variance-based rejection procedure was repeated on epoched channel data, resulting in fewer than 1% rejected single channel trial time series on average (subject range 0.09 – 1.52%).

### Spatial filtering

To emphasize signal reproducibility and across-subject generalization, we applied a spatial filter estimated from single-trial responses to solo speech listening that were included in an independent study (Vanthornhout et al., 2018) using an equivalent EEG system. The filter was obtained through joint decorrelation (de Cheveigné and Parra, 2014; de Cheveigné and Simon, 2008) applied to recordings from 28 subjects listening to a 14.5 minute long story presented by a single talker without noise (Vanthornhout et al., 2018). The purpose of this procedure was to have a continuous, attended stimulus to train the spatial filter across several subjects on an independent reference frame and optimized for inter-subject correlation. To estimate the filter, the independent study recordings were de-meaned, filtered between 1 and 30 Hz with a fourth-order FIR Hamming window filter and corrected for group delay. Two independent components were automatically selected based on their 10-30 Hz region broadband power. Data were downsampled to 64 Hz and, after joint decorrelation, the spatial filter associated with EEG component of the highest evoked/induced activity ratio (de Cheveigné and Simon, 2008) was retained. The4 spatial profile associated with this filter is represented in Figure 2A. All subject data from the present studies were spatially filtered with this fixed single component as a single virtual sensor.

**Figure 2.**
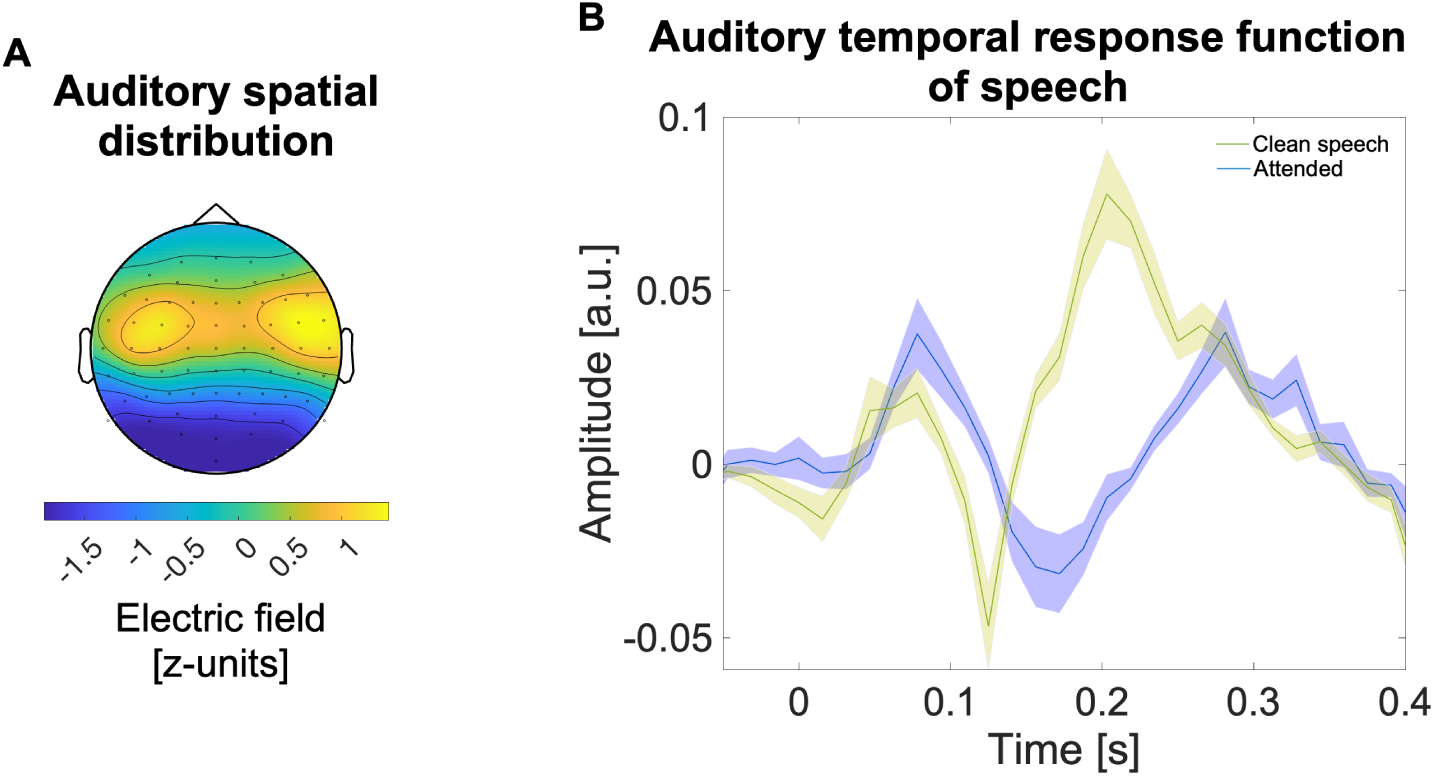
Auditory tracking of the envelope onset signals of solo and CP speech. (**A**). Auditory EEG distribution from the top spatial component resulting from the data-driven spatial filtering procedure applied to EEG data (see Spatial filtering). **(B).** Triphasic auditory temporal response functions (TRFs) during clean and cocktail-party speech listening. Grand averages of TRFs from 71 participants in pre-listening (clean speech condition, yellow) and CP-listening (NP-attended, blue). In pre-listening, participants listen to a clean single speaker, representing optimal conditions for neural speech encoding. By contrast the NP-attended TRF represents processing of a target under the relatively less intelligible cocktail party paradigm. Despite differences in latency, three distinct responses are observed in auditory processing across both paradigms, the positive P1 component as early as 50-75 ms, the negative N1 begins at approximately 125 ms, and the P2 emerges approximately from 200 ms. Across all studies, TRFs model the neural tracking of the envelope onsets of speech.

### Temporal response function

The temporal response function (TRF) is a linear systems method to analyze the neural encoding dynamics in response to changing auditory input over time (Crosse et al., 2016). The stimulus *S*(*t*) was represented by the time series of the continuous speech envelope onset signal, obtained by half-wave rectification of the envelope. Stimulus timeseries involved pre-listening stimuli, *S*_prelistening_(*t*), and the single stream elements of CP stimuli, which was decomposed as *S*_cocktail_(*t*) = *S*_target_(*t*) + *S*_masker_(*t*). In all three cases, *S*_prelistening_(*t*), *S*_target_(*t*), and *S*_masker_(*t*), the timeseries indexes the envelope onsets of an individual (unmixed) speech stream. To consistently represent the execution of the attentional task demands, all trials where participants earned non-positive scores (10-20% of trials) were excluded. For each stimulus attentional category (attended, unattended) and experience condition (AK, UK, NP), single trial stimuli presentations were concatenated and aligned with the corresponding response epochs *r*(*t*), which resulted in data epochs equivalent to up to 15 minutes in pre-listening conditions and 2.5 minutes in cocktail party listening per participant.

The linear model is formulated as:

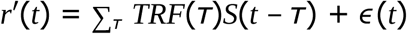

where *r*’(*t*) is the neural response predicted by the TRF model *TRF*(*τ*) under the stimulus representation *S*(*t*), while *ɛ*(*t*) is the residual contribution to the response not explained by the linear model. TRFs were estimated by boosting (David et al., 2007) between concatenated stimulus and EEG time series, scaled to z-units, with 20-fold cross-validation, and analyzed at the –100 to 400 ms window. Each grand average TRF was obtained by averaging the individual TRFs across participants per stimulus attentional category and experience condition.

### Statistical analysis

#### Determination of attentional effect windows

To determine the relevant time windows for attentional selection unbiased by any priming manipulation in the present studies, we determined the timing of attentional effects (AEs) across the cocktail party data. For this, non-parametric tests (Maris, 2011; Martin et al., 2008) were applied using cluster-based non-parametric testing corrected for multiple comparisons. Subject TRFs from the no prior knowledge (NP) condition were used across all three experiments (71 subjects, see Participants sections, below). For each contrast between TRFs to attended versus unattended streams, its *t*-value time series was estimated and thresholded at the *t* distribution’s 5^th^ percentile for randomisation-based testing. Any cluster exceeding the threshold was deemed significant if its associated *t*-statistic (sum of *t*-values within the cluster) exceeded the reference value from the randomization distribution. The distribution was estimated from *N* = 2^17^ resamplings generated by random reshuffling of attended versus unattended conditions per participant. This initial test revealed significant modulations of selective attention on the attended versus unattended speech TRFs, i) between 31-78 ms (corresponding to the P1 window), ii) between 100-180 ms (N1 window), and iii) between 230-320 ms (P2 window), *p* = 1.53×10^−5^. For all three windows, the differences related a higher amplitude for window-averaged TRF estimates to the target than the background speech. The three component windows were then fixed for statistical analyses of TRF data across all experiments.

#### Interaction between selective attention and prior experience

Once the relevant attentional windows were identified at the latencies of the P1, N1 and P2 components (Figure 2B), we examined the statistical interactions between prior experience (AK, UK, and NP conditions) and attention (attended and unattended condition). Two-way repeated measures ANOVAs were performed on time-averaged TRF data as dependent variable, quantified per component and listener. To interpret valid F-ratios, data sphericity violations were determined based on Mauchly testing; for cases where sphericity could not be assumed, Greenhouse-Geisser (ε < 0.75) or Huynh-Feldt (ε > 0.75) corrections were applied (Field, 2017).

#### Classification of negative findings

In case of non-significant effects, Bayesian statistics was additionally performed to determine the strength of evidence for the null hypothesis using the JASP 0.19 package (JASP Team, 2024). The analysis included estimation of the probability of a model M given the data, P(M|data), where M corresponds to either (*i*) a null effect (*H*_0_) model, or a significant effect model of (*ii*) Attention, (*iii*) Experience, (*iv*) Attention + Experience, or (*v*) Attention + Experience + Attention × Experience. To determine which component(s) may be safely excluded as they do not improve predictive performance, P(M| data) of each model was used to quantify the contribution of each participant component. For each component C (Attention, Experience, Attention × Experience), comparisons were performed in two ways: a) corresponding exclusion Bayes Factors (BF_excl_) were calculated as the sum of all p(M without C | data) divided by the sum of all p(M with C | data), i.e. “across all models”; b) restricted to models that only differ in the presence or absence of a given component, i.e. “across matched models”, which provides a conservative estimate (Keysers et al., 2020) of the evidence strength of their null contributions. Therefore we report BF_excl_ as a range, used to classify the strength of evidence of a null finding as anecdotal (1<BF_excl_<=3), moderate (3<BF_excl_<=10), and strong (BF_excl_>10).

### Experiment 1

Experiment 1 – Full priming was designed for participants to gain prior knowledge with a clean auditory stimulus and evaluated the role of this experience in relation to foreground or background speech processing during the decisive selective attention stage. For AK and UK trials, the pre-listening stimulus was therefore identical to one of the two elements of the mixture. In the AK condition, participants were instructed so that they effectively focus, during CP-listening, on a target stream that is identical to the one presented at pre-listening period (Figure 1 shows this case). Conversely, in UK trials, participants were instructed so that they effectively disregard, during CP-listening, a stimulus that is identical with the one presented during the pre-listening stage as it now acts as the background stream. All stimuli were constructed using samples from Stimuli Database 1. The task comprised 108 trials, and the total experimental session lasted approximately 2.5 hours.

### Participants

Thirty-six subjects participated in Experiment 1 (27 female; mean age 26.5 ± 4.7 SD; 7 left-handed). Sample size is determined based on a single-trial auditory TRF study (Cervantes Constantino et al., 2023). One participant was excluded due to a total negative score on the task, while the behavioural response file of the other participant was not saved correctly.

### Stimuli Database 1

Database 1 (D1) consisted of 259 single-speaker audios (129 female voices, mean duration 8.6 s, ±0.8 s). D1 was based on various source formats, such as news broadcasts, audiobooks, and interviews with Uruguayan Spanish speakers. Additional audio clips were extracted from radio and TV channels, podcasts, and YouTube videos. All audio sentences were selected for explicit thematic content, and were issued by a single speaker without background noise. In D1, four keywords were chosen per audio sentence, which included adjectives, nouns, verbs, and adverbs avoiding those present near the end of the sentence. Additionally, one or more topical categories were assigned to each sentence based on fixed pool of database topics that included politics, government, sports, religion, education, health, leisure, personal, culture, nature, labour, commerce, and money. Furthermore, apparent voice gender (male or female) and age (less than or older than 35 years) features were designated according to the source material, resulting in 59 younger and 200 older speakers.

### Task questionnaire and scoring system

The four questionnaire options that involved sentence keywords were randomly drawn from a list of all 8 keywords from the attended and unattended stream. The remaining four options listed possible topics of the target stream, which were presented based on a random selection of four out of 13 database categories. Failure to select any of the 9 options in the questionnaire resulted in a deduction of –2 points (equivalent to missing two correct keywords on average).

## Results

Participants completed the trials with accuracy, i.e., scored 1 point or more, in the no prior knowledge condition (NP, mean success rate 81.0 ± 9.6% SD trials), the attended known condition (AK, 84.3 ± 7.7% trials), and the unattended known condition (UK, 79.0 ± 2.2% trials). A one-way repeated measures ANOVA performed on correct trial rates showed a significant main effect of prior knowledge by full priming (*F*(2, 66) = 5.093, *p* = 0.009). Post-hoc pairwise comparisons revealed a significant difference between AK and UK conditions (corrected *p* = 0.001). Neither the NP vs. UK (corr. *p* = 0.799) nor the AK vs. NP (corr. *p* = 0.267) comparisons reached statistical significance. The results were mirrored in a similar analysis of voice identity based questions, which showed a significant main effect of prior knowledge by full priming (*F*(2, 66) = 6.535, *p* = 0.003). Post-hoc pairwise comparisons of voice identity reports similarly revealed a significant difference between AK and UK conditions (corr. *p* < 0.001). Neither the NP vs. UK (corr. *p* = 0.761) nor the AK vs. NP (corr. *p* = 0.130) comparisons reached statistical significance. In addition, an analysis of lexical accuracy, based on the average number of reported target words less background words, revealed again a significant main effect of prior knowledge by full priming (*F*(2, 66) = 14.358, *p* < 0.001). Post-hoc pairwise comparisons of keyword responses showed a significant difference between AK and UK conditions (corr. *p* < 0.001) and, in addition, between AK and NP conditions (corr. *p* = 0.023). The NP vs. UK comparison did not reach statistical significance (corr. *p* = 0.051). In the case of topic reports, there was no significant effect of prior experience by full priming (*F*(2, 66) = 0.963, *p* = 0.387). Overall, the behavioral data suggest that target priming conferred an advantage on participants’ solving of the cocktail party task questionnaire compared to masker priming; this does not exclude the possibility that background priming conditions may load participants’ working memory relatively more during selection and/or retrieval, leading to weaker performance. For the specific case of lexical accuracy, target priming may in addition facilitate behavior relative to unprimed conditions.

To investigate the statistical interaction between selective attention and prior auditory experience/knowledge of a speech stimulus at each of the relevant speech processing stages (P1, N1, P2), a two-way repeated measures ANOVA was conducted per TRF component estimate of CP listening data.

### P1

A two-way repeated measures ANOVA (Attention × Experience) conducted on TRF estimates of the P1 component did not reveal a statistically significant interaction between selective attention and prior knowledge (*F*(2,66)=0.105, *p*=0.901). At this early stage, there was a significant main effect of selective attention (*F*(1,33)=7.015, *p*=0.012, ρι_p_ =0.175), and there was no significant main effect of prior knowledge (*F*(1.662,54.833)=0.987, *p*=0.366). The data suggest that, while selective attention may modulate the neural tracking of onsets in the envelope of speech at this early stage (Figure 3a), this modulation may not be influenced by prior experience of the target or masker auditory stimuli. Evidence for the absence of the main effect of Experience, and of the Attention × Experience interaction, was assessed through a Bayesian repeated measures ANOVA conducted on the P1 dataset. The exclusion Bayes Factors (BF_excl_) were 5.774-8.235 and 10.061-22.386, respectively, indicating moderate evidence of the absence of the main effect of Experience, and strong evidence of the absence of the Attention × Experience interaction.

**Figure 3.**
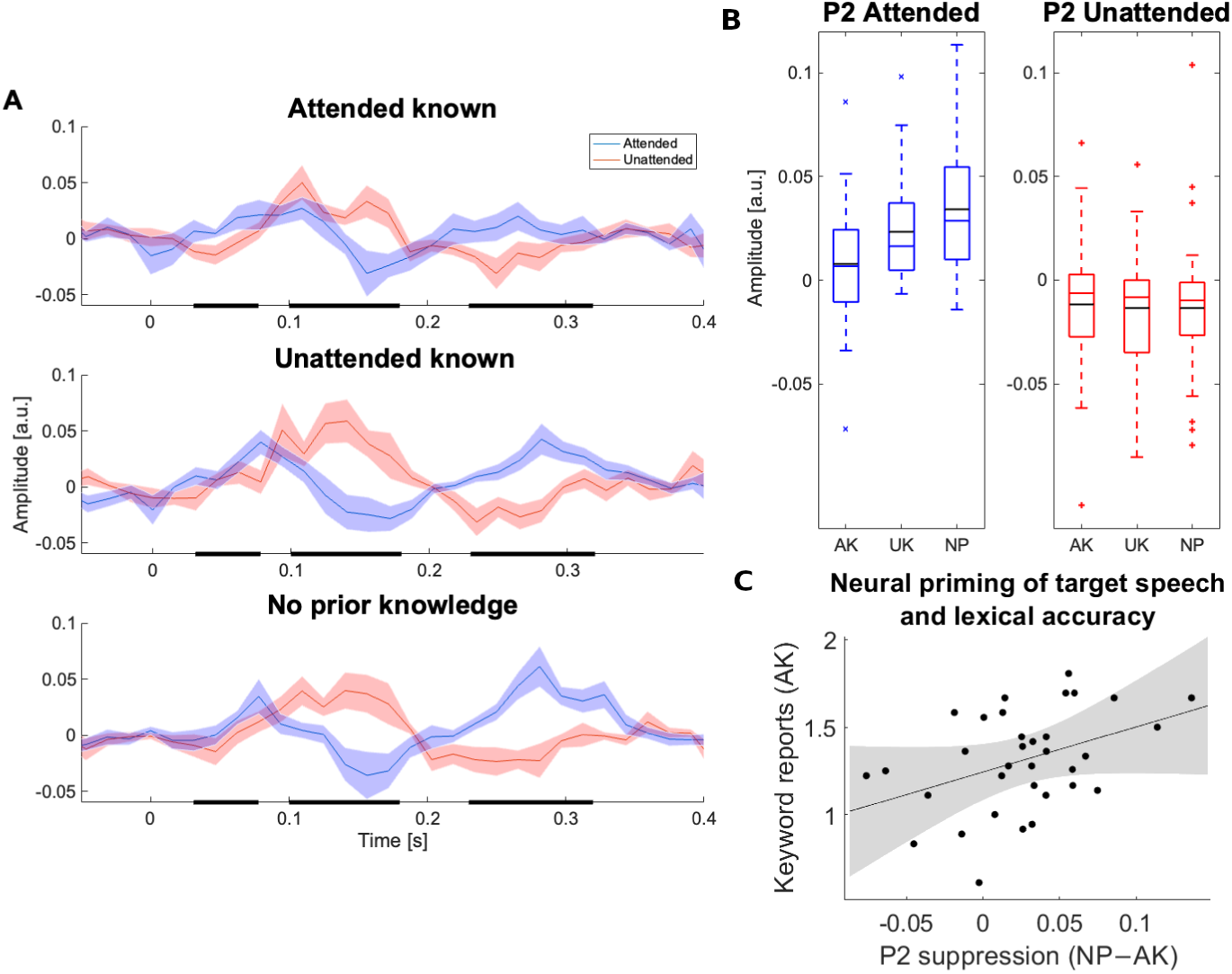
Results from Experiment 1 – Full priming. **(A)** TRF models of neural tracking of the envelope onsets from attended (blue) and unattended speech (red), across prior knowledge conditions. The shaded areas represent the standard error of the mean. The thick bars on the time axis represent analysis windows corresponding to the P1, N1, and P2. **(B)** Interaction between attention and prior knowledge at the P2 stage (230-320 ms). TRF estimates of the P2 from attended and unattended streams under AK, UK and NP conditions, where a significant decrease of the P2 corresponding to the target stream is observed under target priming. **(C)** At the subject level, target priming-led suppression of the P2 component predicts lexical accuracy in target primed trials, which is measured as the average number of target speech words (minus background) reported by participants.

### N1

A two-way repeated measures ANOVA (Attention × Experience) conducted on TRF estimates of the N1 component did not reveal a statistically significant interaction between selective attention and prior knowledge (*F*(2,66)=1.488, *p*=0.233). At this relevant stage, there was a significant main effect of selective attention (*F*(1,33)=26.568, *p*<0.001, ρι_p_ =0.446), and there was no significant main effect of prior knowledge (*F*(2,66)=0.221, *p*=0.802). The data are in agreement with previous findings that selective attention may modulate the neural tracking of onsets in the envelope of speech at this stage, with enhanced responses to the target versus masker speech (Figure 3a). The results however do not support the hypothesis that such modulation is influenced by prior experience of the target or masker auditory stimuli. Evidence for the absence of the main effect of Experience, and of the Attention × Experience interaction, was assessed through a Bayesian repeated measures ANOVA conducted on the N1 dataset. The exclusion Bayes Factors (BF_excl_) were 12.064-13.338 and 2.497-7.922, respectively, indicating strong evidence of the absence of a main effect of Experience, and anecdotal to moderate evidence of the absence of an Attention × Experience interaction.

### P2

A two-way repeated measures ANOVA (Attention × Experience) conducted on TRF estimates of the P2 component revealed a statistically significant interaction between selective attention and prior knowledge (*F*(2,66)=4.745, *p*=0.012, ρι_p_ =0.126). At this late stage, there was a significant main effect of selective attention (*F*(1,33)=49.128, *p*<0.001, ρι_p_ =0.598), and there was no significant main effect of prior knowledge (*F*(2,66)=2.927, *p*=0.061). The data are in agreement with previous findings that selective attention may modulate the neural tracking of onsets in the envelope of speech at this stage, with enhanced responses to the target versus masker speech (Figure 3a). The results furthermore indicate that such modulation is influenced by prior experience of auditory stimuli. Evidence for the absence of the main effect of Experience was assessed through a Bayesian repeated measures ANOVA conducted on the P2 dataset. The exclusion Bayes Factor (BF_excl_) was 0.263-1.660, indicating anecdotal evidence of the absence or presence of a main effect of Experience.

To address the significant interaction, the effect of prior knowledge was evaluated on post-hoc analyses of attended and, separately, unattended speech. A one-way repeated measures ANOVA (Experience) was conducted on TRF estimates of the P2 component related to the attended stream, revealing a significant effect of prior knowledge (*F*(2,66)=7.186, corr. *p*=0.004). A similar analysis conducted on TRF estimates related to the unattended stream did not reveal a significant effect of prior knowledge at the P2 (*F*(2,66)=0.044, corr. *p*>0.957). For this case, the corresponding exclusion Bayes Factor (BF_excl_) was 10.862, indicating strong evidence of the negative finding consistent with the significant statistical interaction. Hence, priming modulated the neural coding of envelope onsets from the attended stream, while for unattended streams these responses remained similar across priming conditions.

Subsequent post-hoc pairwise comparisons of TRF estimates of the attended stream at the P2 stage revealed a significant difference between AK and NP conditions (corr. *p* = 0.006). As a result of target priming, a reduced P2 was observed for known targets in comparison to control conditions. Neither the NP vs. UK (corr. p = 0.289) nor the AK vs. UK (corr. p = 0.082) comparisons reached statistical significance (Figure 3b).

Having observed the P2 interaction explained in terms of a AK-led reduction, we subsequently explored whether inter-subject differences, taken as indicators of neural priming of the speech targets at the individual level, demonstrate a behavioral correlate. Our initial test did not evidence that these neural measures were linearly correlated with subjects’ percent correct trials (Pearson’s *r*=-0.075, *p*=0.674), which were scored based on specific dimensions of speech segregation, including accurate target voice and topic identification (see *Methods*). We inspected the sub-dimension of lexical accuracy, measured in terms of average correct (target) minus incorrect (masker) reported words, since it represented the most appropriate behavioral dimension resulting from precise temporal segmentation of the target stream. This exploratory approach suggests that, at the individual level, participants that show greatest neural priming related to target speech in a cocktail party are also the most able to report target words, evidenced by a significant linear correlation (*r*=0.411, *p*=0.016, bootstrapped CI [0.156 – 0.601]; Figure 3c). To validate the reliability of individual observations, a jackknife procedure was performed on the linear correlation analyses, showing significant positive associations in all resamplings. The significant finding was confirmed as robust to potential outliers, as indicated by robust linear regression with Huber weighting (*t*(32)=2.275, *p*=0.021).

In summary, the results confirm cocktail party paradigm findings that selective attention improves the neural tracking of envelope onsets from attended speech compared to unattended speech at the P2 stage. This study shows that such attentional enhancement can be modulated by prior experience. Changes consist of reduced tracking of envelope onsets from attended speech when the observer has gained expectations about the same single auditory stimulus through exact priming.

### Experiment 2a

Experiment 2a – Voice priming was designed for participants to gain prior knowledge that is limited to a speaker’s voice, and evaluated the role of this experience in relation to foreground or background speech processing during the decisive selective attention stage. NP trials were identically generated as in Experiment 1. AK/UK trials were specifically generated so that each new speaker presenting at pre-listening corresponded with one of the two speakers of the mixture during the cocktail party presentation. In the AK condition, participants were instructed to focus on a target stream whose speaker’s voice was in effect the same as presented during the pre-listening period, thus with voice-priming of the target. Conversely, in UK trials participants were instructed so as to effectively disregard a speaker whose voice coincided with that presented during the pre-listening stage, thus with voice-priming of the masker. Repeating speaker voice stimuli pairs always involved different sentences and contents. AK/UK stimuli were exclusively sampled and constructed from Stimuli Database 2 (see below). We discarded the possibility that same content sentence pairs (including paraphrases) were used at any point for a given participant. The experiment consisted of 78 trials and the total duration of the experimental session was approximately 1.5 hours.

### Participants

Seventeen subjects participated in Experiment 2a (7 female; mean age 24.1 ±4.8 SD; 3 left-handed). In our repeated measures F-test design (1 group, 6 measurements), replicating the smallest significant effect observed in Experiment 1 (ρι ^2^=0.126, equivalently, Cohen’s f=0.38) would suggest a minimum sample size of 9 for 80% statistical power or 17 for 99% statistical power.

### Stimuli Database 2

Stimuli Database 2 (D2) consisted of 348 single-speaker audios containing 184 female voices. The mean duration of the audios was 8.08 s, SD ± 1.97 s. Individual sentences were created and recorded using the Microsoft Research Paraphrase Corpus (MRPC, Quirk et al., 2004), a sentence database that contains phrase pairs with similar information about international news from different newspapers and sources. We selected 169 paraphrase pairs that were rated by the MRPC as semantically equivalent, and translated them into Spanish. Non-Spanish names (e.g., people, companies, or localities) in these sentences were substituted with Spanish, familiar, local or regional names using an online random name generator (https://generadordenombres.online/). An example of a resulting paraphrase pair used for D2 audios is shown in Table 2.

**Table 1.** Example of Stimuli Database 1 voice and sentence annotation entries used for questionnaire testing.

| # | Politics | Gov't | Sports | Religion | Educ. | Keyw1 | Keyw2 | Keyw3 | Keyw4 | M/F | Age |
| --- | --- | --- | --- | --- | --- | --- | --- | --- | --- | --- | --- |
| 1 | X | X |  | X |  | Biases | Brain | Financial | Invest | F | Younger |
| 2 | X |  |  |  |  | Retire | Write | Journal | Stuff | M | Older |
| 3 |  | X |  | X |  | Tourism | Illusion | Transform | Experience | M | Younger |
| 4 | X |  |  |  |  | Advances | Rights | Money | Depression | F | Older |

**Table 2.** Example of a paraphrase pair used in Stimuli Database 2.

| MRPC sentence 202294 | MRPC sentence 202143 (paraphrase) |
| --- | --- |
| Original: *The proceedings were taken up with prosecutors outlining their case against Amrozi, reading 33 pages of documents outlining allegations against him.* | Original: *Proceedings were taken up with prosecutors outlining their case against Amrozi, reading a 33-page accusation letter to the court.* |
| Translation: [REMOVED] | Translation: [REMOVED] |

Audio recordings were obtained from local volunteers who read aloud one or two pairs of paraphrases as simulating a spontaneous conversation. An Olympus recorder (VN-541OC, OM Digital Solutions Corporation, Japan) was used with settings suitable for recording in the open air, at 44.1 kHz sampling rate, 40 Hz – 13 kHz frequency response and 5 – 320 kbps bit rate range. Selected recordings were made in noise-free open or closed spaces, and contained clear verbatim message narrations of the indicated sentence(s). Resulting audios were annotated for their speaker age, sex and keywords as in D1, with the exception that keywords were limited to two per phrase. The annotated keywords always differed across paraphrasing pairs.

### Task questionnaire and scoring system

The four questionnaire options involving sentence keywords were randomly drawn from a list of 4 keywords built from the attended and unattended sentences. The remaining four options listed possible topics of the target stream, which were presented based on a random selection of four out of 14 topic categories in D2. Failure to select any of the 9 options in the questionnaire resulted in a score of 0 for the trial. In some participants, it was possible that a limited number of trials (≤18) may show keywords from the unattended (background) sentence only; these trials were not considered for the effects of lexical accuracy analyses.

## Results

Participants completed the trials with accuracy in the NP condition (87.6 ± 9.0% correct trials), in the AK condition (80.1 ± 15.3% trials), and in the UK condition (87.8 ± 10.3% trials). A one-way repeated measures ANOVA performed on correct trial rates did not show a significant main effect of prior knowledge by voice priming (*F*(1.151, 18.422) = 4.020, *p* = 0.055). A similar analysis of voice identity based questions likewise did not show a significant main effect of prior knowledge by voice priming (*F*(2, 32) = 1.108, *p* = 0.343). In addition, an analysis of lexical accuracy did not reveal a significant main effect of prior knowledge by voice priming (*F*(1.277, 20.425) = 1.240, *p* = 0.292). In the case of topic reports, there was no significant effect of prior experience by voice priming (*F*(2, 32) = 0.005, *p* = 0.995).

To investigate the statistical interaction between selective attention and prior auditory experience/knowledge of a speaker’s voice at each of the relevant speech processing stages (P1, N1, P2), a two-way repeated measures ANOVA was conducted per TRF component estimate of CP listening data.

### P1

A two-way repeated measures ANOVA (Attention × Experience) conducted on TRF estimates of the P1 component did not reveal a statistically significant interaction between selective attention and prior knowledge (*F*(2,32)=0.276, *p*=0.761). At this early stage, there was no significant main effect of selective attention (*F*(1,16)=4.471, *p*=0.051), and there was no significant main effect of prior knowledge (*F*(2,32)=1.577, *p*=0.222). The data from Experiment 2a do not confirm the finding from Experiment 1 that selective attention may modulate the neural tracking of onsets in the envelope of speech at this early stage. Similar to Experiment 1, this component was not found to be influenced by prior experience of the target or masker speaker’s voice. Evidence for the absence of the main effects of Experience, Attention, and of the Attention × Experience interaction, was assessed through a Bayesian repeated measures ANOVA conducted on the P1 dataset. The exclusion Bayes Factors (BF_excl_) were 3.568-4.772, 0.951-1.358, and 4.873-10.770, respectively, indicating moderate evidence of the absence of the main effect of Experience, anecdotal evidence of the absence or presence of the main effect of Attention, and moderate to strong evidence of the absence of the Attention × Experience interaction.

### N1

A two-way repeated measures ANOVA (Attention × Experience) conducted on TRF estimates of the N1 component did not reveal a statistically significant interaction between selective attention and prior knowledge (*F*(2,32)=0.029, *p*=0.971). At this relevant stage, there was a significant main effect of selective attention (*F*(1,16)=18.670, *p*=0.001, ρι_p_ =0.539), and there was no significant main effect of prior knowledge (*F*(1.438,23.001)=2.187, *p*=0.146). As in Experiment 1, the data are in agreement with previous findings of auditory object segregation via enhanced responses to foreground versus background speech at this stage (Figure 4). Similarly, the results do not support the hypothesis that such modulation is influenced by prior experience of the target or masker speaker’s voice. Evidence for the absence of the main effect of Experience, and of the Attention × Experience interaction, was assessed through a Bayesian repeated measures ANOVA conducted on the N1 dataset. The exclusion Bayes Factors (BF_excl_) were 1.286-1.539 and 3.565-6.792, respectively, indicating anecdotal evidence of the absence of a main effect of Experience, and moderate evidence of the absence of an Attention × Experience interaction.

**Figure 4.**
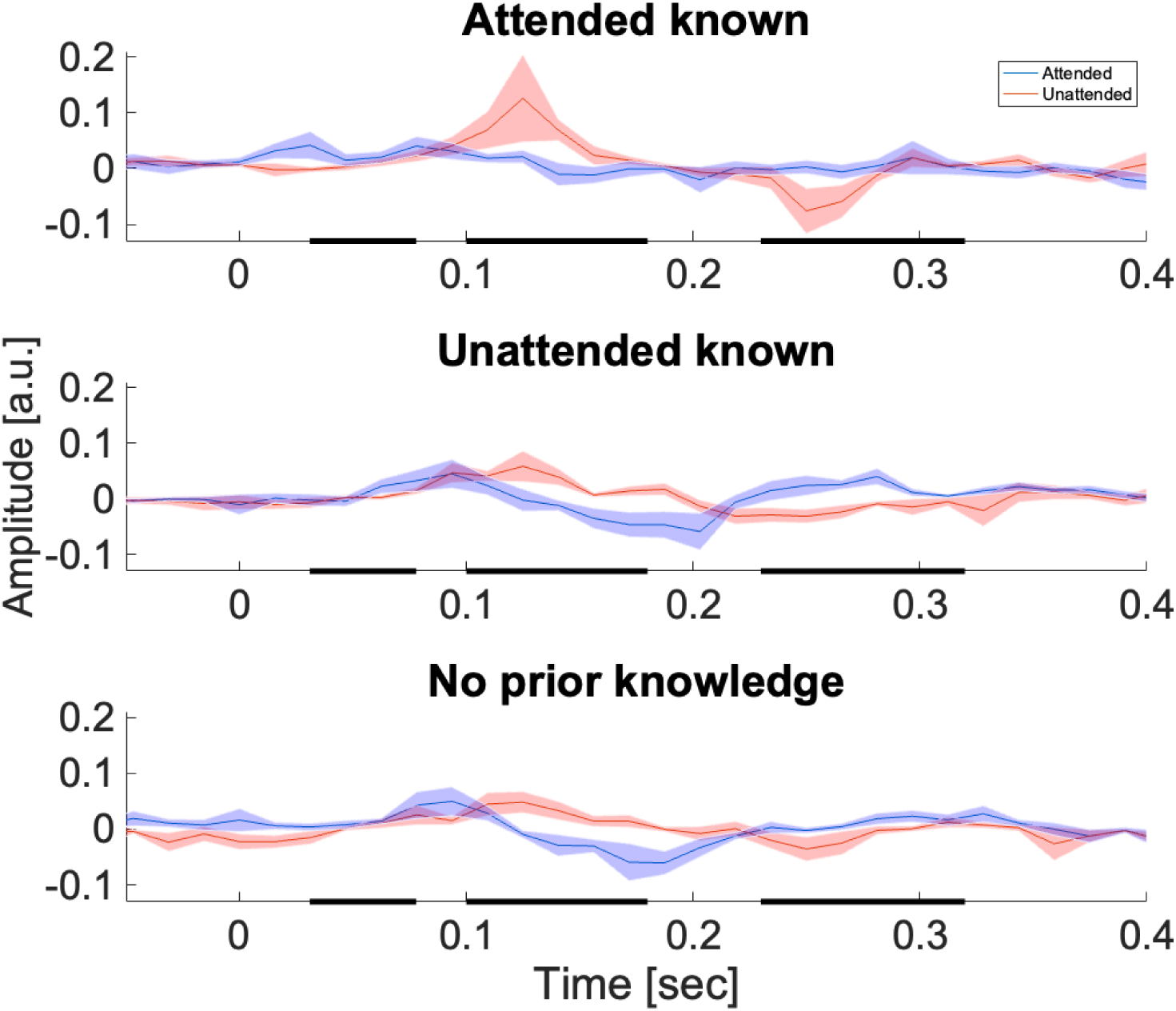
TRF results from Experiment 2A – Voice priming. TRF models of neural tracking of the envelope onsets from attended (blue) and unattended speech (red), across speaker’s voice prior knowledge conditions. The shaded areas represent the standard error of the mean. The thick bars on the time axis represent analysis windows corresponding to the P1, N1, and P2.

### P2

A two-way repeated measures ANOVA (Attention × Experience) conducted on TRF estimates of the P2 component did not reveal a statistically significant interaction between selective attention and prior knowledge (*F*(2,32)=1.053, *p*=0.361). At this late stage, there was a significant main effect of selective attention (*F*(1,16)=19.206, *p<*0.001, ρι_p_ =0.546), and there was no significant main effect of prior knowledge (*F*(1.310, 20.957)=0.638, *p*=0.474). The data are also in agreement with previous findings that selective attention may modulate the neural tracking of onsets in the envelope of speech at this stage, with enhanced responses to the target versus masker speech (Figure 4). The results however do not support the hypothesis that such modulation is influenced by prior experience of the target or masker speaker’s voice. Evidence for the absence of the main effect of Experience, and of the Attention × Experience interaction, was assessed through a Bayesian repeated measures ANOVA conducted on the P2 dataset. The exclusion Bayes Factors (BF_excl_) were 3.905-4.129 and 2.698-3.365, respectively, indicating moderate evidence of the absence of a main effect of Experience, and anecdotal evidence of the absence of an Attention × Experience interaction.

### Experiment 2b

Experiment 2b – Content priming was designed for participants to gain prior knowledge that is limited to a spoken message’s content, and evaluated the role of this experience in relation to foreground or background speech processing during the decisive selective attention stage. NP trials were identically generated as in the previous experiments. AK/UK trials were specifically generated so that each new spoken message presented at pre-listening corresponded with one of the two spoken messages of the mixture during the cocktail party presentation. In the AK condition, participants were instructed to focus on a target stream whose spoken content was in effect the same as presented during the pre-listening period, thus with content-priming of the target. Conversely, in UK trials participants were instructed so as to effectively disregard a stream whose content coincided with that presented during the pre-listening stage, thus with content-priming of the masker (see Table 3 for an example of AK and UK trial sentence combinations). In this experiment, all speaker voices always differed from each other, within and across trials. As in Experiment 2a, AK/UK stimuli were exclusively sampled and constructed from D2. The presentation order of the paraphrase sentence pairs was equally counterbalanced across participants. The task consisted of 78 trials, and the experimental session lasted approximately 1 hour and 30 minutes.

**Table 3.**
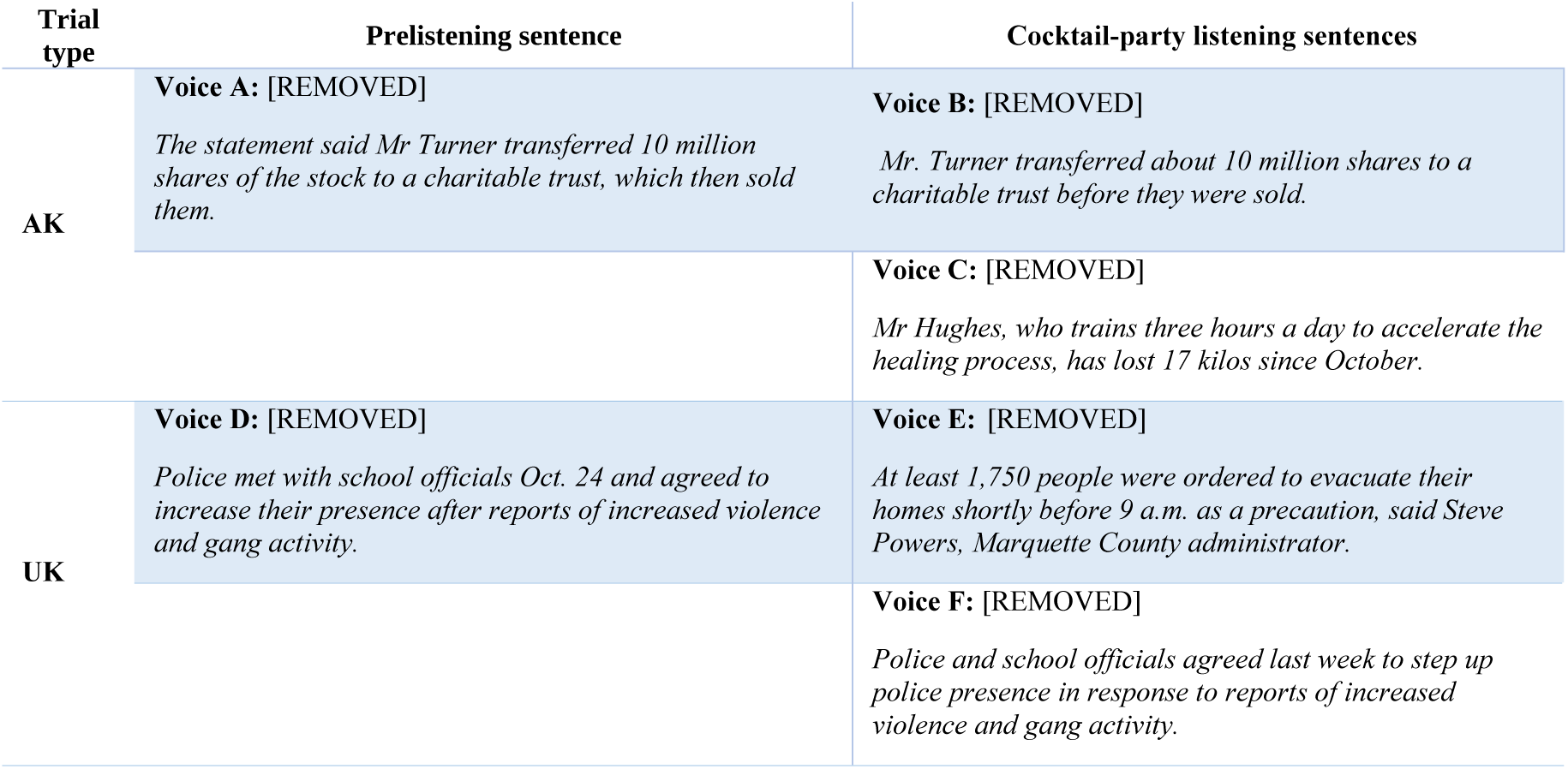
Schematic of a sentence combination for content priming trials in Experiment 2b, illustrating AK and UK conditions, along with original MRPC versions in English. Highlight indicates the instructed stream to attend.

### Participants

Twenty-one subjects participated in Experiment 2b (16 female; mean age 25,7 ± 6 SD; 1 left-handed).

## Results

Participants completed the trials with accuracy in the NP condition (90.1 ± 8.3% correct trials), in the AK condition (84.4 ± 12.7% trials), and in the UK condition (86.2 ± 7.7% trials). A one-way repeated measures ANOVA performed on correct trial rates did not show a significant main effect of prior knowledge by content priming (*F*(2, 40) = 1.921, *p* = 0.160). A similar analysis of voice identity based questions likewise did not show a significant main effect of prior knowledge by content priming (*F*(2, 40) = 1.787, *p* = 0.181). In addition, an analysis of lexical accuracy did not reveal a significant main effect of prior knowledge by content priming (*F*(2, 40) = 1.735, *p* = 0.189). In the case of topic reports, there was no significant effect of prior experience by content priming (*F*(2, 40) = 2.332, *p* = 0.110).

To investigate the statistical interaction between selective attention and prior auditory experience/knowledge of voiced content at each of the relevant speech processing stages (P1, N1, P2), a two-way repeated measures ANOVA was conducted per TRF component estimate of CP listening data.

### P1

A two-way repeated measures ANOVA (Attention × Experience) conducted on TRF estimates of the P1 component did not reveal a statistically significant interaction between selective attention and prior knowledge (*F*(2,40)=0.201, *p*=0.819). At this early stage, there was a significant main effect of selective attention (*F*(1,20)=8.982, *p*=0.007, ρι ^2^=0.310), and there was no significant main effect of prior knowledge (*F*(2,40)=0.904, *p*=0.413). The data from Experiment 2b replicate the finding from Experiment 1 that selective attention may modulate the neural tracking of onsets in the envelope of speech at this early stage. Similar to Experiments 1 and 2a, this component was not found to be influenced by prior experience of the target or masker speech content (Figure 5). Evidence for the absence of the main effect of Experience, and of the Attention × Experience interaction, was assessed through a Bayesian repeated measures ANOVA conducted on the P1 dataset. The exclusion Bayes Factors (BF_excl_) were 4.262-5.735 and 6.280-9.922, respectively, indicating moderate evidence of both absence findings.

**Figure 5.**
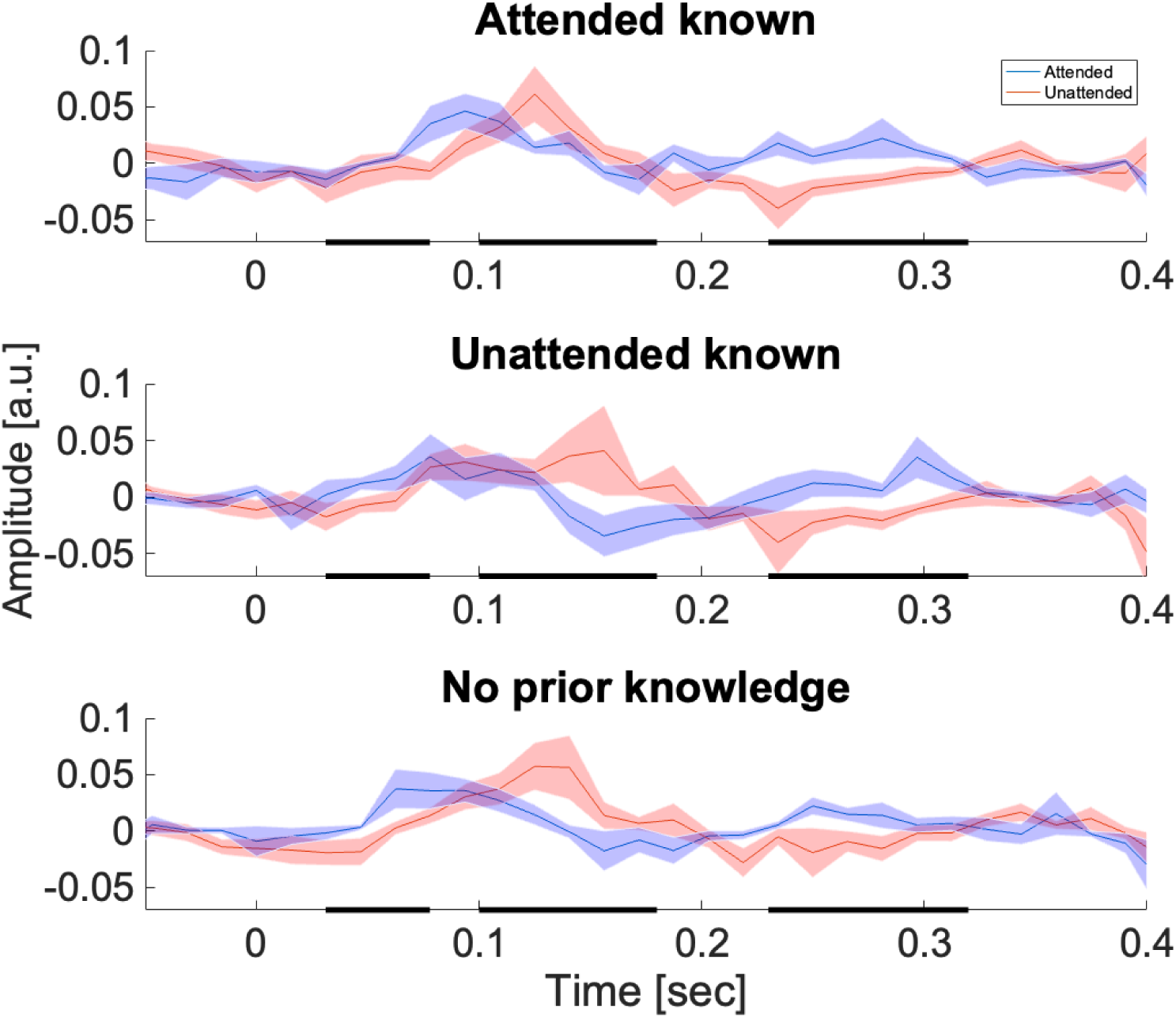
TRF results from Experiment 2b – Content priming. TRF models of neural tracking of the envelope onsets from attended (blue) and unattended speech (red), across speech content prior knowledge conditions. The shaded areas represent the standard error of the mean. The thick bars on the time axis represent analysis windows corresponding to the P1, N1, and P2.

### N1

A two-way repeated measures ANOVA (Attention × Experience) conducted on TRF estimates of the N1 component did not reveal a statistically significant interaction between selective attention and prior knowledge (*F*(2,40)=1.122, *p*=0.336). At this relevant stage, there was a significant main effect of selective attention (*F*(1,20)=8.025, *p*=0.010, ρι_p_ =0.286), and there was no significant main effect of prior knowledge (*F*(2,40)=0.663, *p*=0.521). As in Experiments 1 and 2a, the data are in agreement with previous findings of auditory object segregation via enhanced responses to foreground versus background speech at this stage (Figure 5). Likewise, the results do not support the hypothesis that such modulation is influenced by prior experience of the target or masker speech content. Evidence for the absence of the main effect of Experience, and of the Attention × Experience interaction, was assessed through a Bayesian repeated measures ANOVA conducted on the N1 dataset. The exclusion Bayes Factors (BF_excl_) were 6.844-7.568 and 2.197-5.593, respectively, indicating moderate evidence of the absence of a main effect of Experience, and anecdotal to moderate evidence of the absence of an Attention × Experience interaction.

### P2

A two-way repeated measures ANOVA (Attention × Experience) conducted on TRF estimates of the P2 component did not reveal a statistically significant interaction between selective attention and prior knowledge (*F*(2,40)=0.664, *p*=0.520). At this late stage, there was a significant main effect of selective attention (*F*(1,20)=21.299, *p*<0.001, ρι_p_ =0.516), and there was no significant main effect of prior knowledge (*F*(2,40)=0.352, p=0.706, Figure 7). The data confirm our previous experiment findings that selective attention may modulate the neural tracking of onsets in the envelope of speech at this stage with enhanced responses to the target versus masker speech. Similarly to partial priming results in Experiment 2a, the results however do not support the hypothesis that such attentional modulation is influenced by prior experience limited to the target or masker speech content. Evidence for the absence of the main effect of Experience, and of the Attention × Experience interaction, was assessed through a Bayesian repeated measures ANOVA conducted on the P2 dataset. The exclusion Bayes Factors (BF_excl_) were 6.283-7.168 and 4.468-7.815, respectively, indicating moderate evidence of both absence findings.

## Summary results

Envelope onset coding across studies showed an interaction between selective attention and prior knowledge that was limited to the P2 component, and contingent on exact (full) priming conditions. No interactions with or main effects of prior experience were observed at any other response component or condition (Table 4). With regards to attentional effects, envelope onset coding at the N1 and P2 components were systematically modulated by attention, consistent with previous studies. The results provide additional partial evidence of the attentional modulation of the P1 component.

**Table 4.** Summary of neural results listing statistical significance p-values for interactions and main effects across components and experiments, during speech envelope onset EEG tracking. Bold entries indicate significant effects. Letters indicate classification of negative findings evidence by BF_excl_. a, anecdotal, <3; m, moderate, 3 to 10; s, strong, >10. (+) indicates evidence for the presence of the effect (Bf_excl_<1). Effect sizes (ρι ^2^ values) are in parentheses.

| Experiment | Effect | P1 (31-78 ms) | N1 (100-180 ms) | P2 (230-320 ms) |
| --- | --- | --- | --- | --- |
| 1 – Full priming | Attention | **0.012 (0.175)** | **<0.001 (0.446)** | **<0.001 (0.598)** |
|  | Experience | 0.366<sup>m</sup> (0.029) | 0.802<sup>s</sup> (0.007) | 0.061<sup>a-a(+)</sup> (0.081) |
|  | Attention x Experience | 0.901<sup>s</sup> (0.003) | 0.233<sup>a-m</sup> (0.043) | **0.012 (0.126)** |
| 2a – Voice priming | Attention | 0.051<sup>a-a(+)</sup> (0.218) | **0.001 (0.539)** | **<0.001 (0.546)** |
|  | Experience | 0.222<sup>m</sup> (0.090) | 0.146<sup>a</sup> (0.120) | 0.474<sup>m</sup> (0.038) |
|  | Attention x Experience | 0.761<sup>m-s</sup> (0.017) | 0.971<sup>m</sup> (0.002) | 0.361<sup>a-m</sup> (0.062) |
| 2b – Content priming | Attention | **0.007 (0.310)** | **0.010 (0.286)** | **<0.001 (0.516)** |
|  | Experience | 0.413<sup>m</sup> (0.043) | 0.521<sup>m</sup> (0.032) | 0.706<sup>m</sup> (0.017) |
|  | Attention x Experience | 0.819<sup>m</sup> (0.010) | 0.336<sup>a-m</sup> (0.053) | 0.520<sup>m</sup> (0.032) |

## Discussion

Our results demonstrate the interaction between selective attention and prior experience during the neural processing of key temporal edge signals from continuous natural speech. This interaction is evident through the suppressed neural tracking response to envelope onsets when predictable speech stimuli exactly match the streamed target within a ‘cocktail-party’ mixture. The decrement, specific to the P2 component of this temporal landmark-locked response, did not revert the attentional gain that is observed at this stage. Priming only partial stimulus information, such as the speaker’s voice, or its speech content alone, by contrast, did not lead to similar changes during cocktail party listening.

An immediate goal of top-down auditory selection is to infer and connect every temporally separate sound element that is caused by the same target source. In such tasks, onset edges represent crucial cues for the temporal segmentation process of listening to masked speech. To understand the mechanisms of their selective tracking adjusted by the observer’s expectations, we manipulated exposure to target or masker speech elements before listeners operated selective attention within speech mixtures. Our single-trial results are consistent with the notion that, during selection, the neural P2 stage (250-300 ms latency) effectively represents the neurally denoised template of the listener’s target speech object. This was supported by findings of fewer processing of its temporal edges when these were exactly predictable from short-term prior experience. Preliminary findings that accurate behavior reporting target speech words may be predicted by the neural reduction lend additional support to the hypothesis that the P2 contains prediction error about the temporal structure of target speech amid masking. Placing apart all relevant listening timepoints through informationally masked scenarios, the brain takes into late account the observer’s precise model of when they expect to listen to the most reliable sensory evidence.

### Experience-related changes on the selective auditory processing of speech after the key N1 stage

*P1.* Cortical encoding of transient increases in the speech envelope may exhibit at least three well-differentiated components in scalp signals (<u>Fiedler et al., 2019</u>; <u>Gansonre et al., 2018, Kraus et al., 2021</u>; <u>Wikman, et al., 2024, Peterson et al., 2017, Han, 2010</u>). The first of them, the P1, appears as a relatively small deflection peaking within 50-70 ms of the impulse encoding response. Convergent findings from ERP and TRF studies suggest that its generators lie within Heschl’s gyrus (<u>Aiken and Picton, 2008</u>; <u>Brodbeck et al., 2020</u>) with potential contributions from the hippocampus, posterior planum temporale, and the lateral temporal cortex (<u>Han, 2010</u>). The P1 stage has been often linked to bottom-up responses from excitatory activity coding acoustically defined stimulus features (Grunwald et al., 2003; Lewald and Getzmann, 2015), e.g., timbre, intensity, or location features of speech. Compared to subsequent scalp components, this early signal appears of a weaker magnitude and is less consistently featured across studies of speech tracking. Our global analysis showed that the P1 is already modulated by top-down auditory selective attention during the cocktail party, a finding that was replicated in some but not all experiments. Such preliminary results appear consistent with previous work showing that top-down goal-directed attention may modulate early latency sensory responses (<u>Golubic et al., 2019</u>; <u>Jones et al., 2016; Luck et al., 1990</u>).

Regarding experience, no effects or interactions with selective attention were observed at the P1. P1 responses to sequences of identical auditory stimuli may undergo repetition suppression effects that have been previously interpreted as automatic filtering out redundant sensory information, or ‘gating’ (<u>Haykin and Chen, 2005</u>; <u>Jones et al., 2016</u>). Such effects, however, appear dissociable from those that emerge as a result of stimuli matching higher-order probabilistic contingencies about auditory sequences (i.e., expectation suppression) and which may be observable at later latencies (Todorovic and de Lange, 2012). As no P1 reductions were observed under any individually repeated target or masker streams within the mixture (which was strictly novel), our results appear consistent with this stage representing speech input with both streams’ temporal structures combined. The finding of a task-related selective modulation may separate these early filtering or gating phenomena from early, partial segregation biases hypothetically linked to passive filtering mechanisms (<u>Brodbeck et al., 2020</u>) under voluntary attention.

*N1.* The second relevant cortical auditory component, the N1, usually appears 100-150 ms post stimulus onset and is sometimes used as a reference point to distinguish early from late selection (<u>Fiedler et al., 2019</u>; <u>Giard et al., 2000</u>). The N1 response to the speech envelope is consistently modulated by top-down attention, which underscores its role in task-oriented stream segregation (Ding and Simon, 2012; Zion Golumbic et al., 2013; <u>Brodbeck et al., 2020</u>; <u>Fiedler et al., 2019</u>). Because by this stage responses reflect parallel, object-based processing of the expected target amid its scene, responses therein have been hypothesized to correspond to the active process of auditory object formation and segregation (<u>Ding and Simon, 2012</u>; <u>Kaya and Elhilali, 2017</u>). This refers to the grouping of sound mixture elements as temporally connected objects, segregated from their scene through extraction of relevant acoustic features, and the binding of sensory evidence likely matching the target object features’ common cause (<u>Bronkhorst, 2000</u>; <u>Moore and Gockel, 2012</u>; <u>Middlebrooks and Simon, 2017</u>; <u>Shinn-Cunningham et al., 2017</u>; <u>Brodbeck et al., 2020</u>; <u>Giraud and Poeppel, 2012</u>; <u>Luo and Poeppel, 2007</u>; <u>Obleser and Kayser, 2019</u>; <u>Ozmeral et al., 2021</u>; <u>Poeppel and Assaneo, 2020</u>; <u>Shinn-Cunningham et al., 2017</u>). One critical cue to extract from voiced signals is their envelope onset, since it corresponds to the temporal sequence that naturally frames segmentation of connected speech (<u>Hertrich et al., 2012</u>). Under selective attention, N1 responses to envelope onset cues extracted from task-relevant speech are similarly subject to gain effects and, in addition, to expedited processing (<u>Brodbeck et al., 2020</u>). Both phenomena underscore the computational role of temporal edge extraction for object formation and segregation under informational masking. Attenuation and delay of edge cues associated with the background stream moreover reflect the gradual establishment of noise-invariant neural representations of a target in a scene as processing advances from the periphery through the auditory cortex (<u>Rabinowitz et al., 2013</u>).

In our results, N1 attentional gains did not show a modulation by any of our priming conditions, as listener expectations about individual scene objects failed to either sharpen or decrease selective tuning at this stage. Previous evidence from auditory studies is mixed with reports of decreases, increases, or null effects on attentional gains under stimulus predictability at the N1 (see reviewed in Lange, 2013). Primarily based on single-tone processing paradigms, this prior work suggests that low-level auditory expectations in some cases influence attentional signals earlier than the auditory responses to dual-stream, continuous speech observed here. Attentional signals are hypothesized to stem from consistent synchronization of endogenous neural activity to sub-elements of the mixed temporal structure that accord with the likely target (<u>Kachlicka et al., 2022</u>). Such a gain system necessarily involves the additional synchronizing of endogenous activity to that background elements, possibly to a lesser degree of consistency, or at a different phase (Lakatos et al., 2013), for instance. One reason for this is that effective discrimination relies on inference and labeling of temporal target groups from the unfolding mixed (or partially unmixed) signal. In the present scenario, this acoustic mixed signal contains auditory uncertainty not explained by prior experience of an individual stream. We propose that the N1 stage overlaps with a putative inference and labeling process undertaken to effectively minimize the net auditory uncertainty at hand. This is consistent with N1 synchronization gains reflecting a dual neural representation, preceding and distinct to that whence distractor elimination reveals the neurally segregated target. The hypothetical post-N1 subordinate representation is then defined by minimal masker-related uncertainty. This predicts that from this point, further reductions in auditory uncertainty primarily relate to the target. Reductions may be obtained via advance templates of the segregated object held in the form of observer temporal expectations from working memory, for instance.

*P2.* The P2 component appears as a prominent positive peak (<u>Alain and Tremblay, 2007, Wang et al., 2020, Verschueren et al., 20</u>20) typically beginning 150 to 200 ms post stimulus onset, or later in aging populations (<u>Pinto et al., 2019</u>; <u>Lewald and Getzmann, 2015</u>; <u>Getzmann et al., 2016</u>). Consistent with previous work (<u>Charest et al., 2009; Brodbeck et al., 2020</u>) our P2 results demonstrated attentional gain effects throughout our experimental manipulations, with limited or no tracking responses to the masker (<u>Fiedler et al., 2019</u>). Activity generating the P2 component has been considered in relation to the encoding of the neurally segregated target (<u>Broderick et al., 202</u>2) involving working memory to extract higher-order semantic, lexical, and contextual information of individual auditory streams. The high attentional selectivity exhibited by this component is modulated by memory demands and behavioral relevance of the task (<u>O’Sullivan et al., 2015</u>). In visual selection, attentional gains of neural responses at this latency are a predictor of working memory performance (<u>Gazzaley and Nobre, 2012</u>). For speech processing, by this stage the neural representations of the masker may no longer be accessible to working memory (<u>Power et al., 2012</u>), although evidence appears mixed (<u>Hjortkjær et al., 2020</u>). Critically, the P2 was sensitive to prior experience of the exact target sequence, a finding that we next discuss in detail.

### Exact priming manipulations induce experience neural changes in selective speech listening

Prior experience with voiced speech enables the brain to establish predictions that may influence auditory processing (<u>Daly and Pitt, 2023</u>; <u>Pinto et al., 2019</u>). Our full priming study (Experiment 1) demonstrated the interaction between selective attention and prior experience at the P2 encoding stage. Specifically, exact foreknowledge of the target stream led to reduced neural processing of its temporal patterning amid masker speech. The suppression of a repeated scene sub-element with a coincident temporal pattern as the primer is consistent with research showing that precise expectations about the temporal occurrence of speech sounds may lead to reduced activity (<u>Pinto et al., 2019</u>). In predictive processing of speech, coincidence of an incoming input pattern with internal expectations reduces the discrepancy or prediction error that is considered to drive, in part, neural tracking activity (<u>Broderick et al., 2019</u>; <u>Park et al., 2023</u>). According to this predictive coding account, cortical activity showing repetition suppression effects might be mediated by supragranular prediction error units, weakening progressively as repetitions reconcile input signal activity with predictions from descending projections (Auksztulewicz and Friston, 2016). Alternative mechanisms may include diminished neural populations involved in the coding of expected stimuli, or populations of a similar size but which operate on shorter processing timeframes, or a combination of these (Grill-Spector et al., 2006). Importantly, as our results show, suppressive effects appear to be contingent on the deployment of attention and diminish or are absent for ignored stimuli (Larsson and Smith, 2012). Our results suggest that attenuation corresponds to the adaptive parsing of target segmentation cues, in this case through prior learning of its temporal structure.

In a MEG study, <u>Wang et al.</u> (<u>2019</u>) investigated the effect of priming target speech on envelope-tracking responses in the cocktail party. Following a multi-trial design, observers were either primed with the exact target speech sequence or, alternatively, they were presented with no advance solo speech, and responses within cocktail party mixtures were analyzed. In primed conditions, neural encoding of the speech envelope was modulated by attention around 100 ms; in unprimed trials, this modulation appeared unclear, however. Source analyses revealed that activity changes related to the differential attentional effect by priming were localized at the bilateral superior temporal sulcus (STS) (<u>Wang et al., 2019</u>). While the interaction between attention and priming was not specifically addressed at individual TRF components in their study, across analysis windows (50-600 ms) the results appeared consistent with target priming suppressing the representation of masker (rather than target) speech. Discrepancies with our results may relate to the experimental design of both tasks, which in their study addressed acoustic gap detection of a target stream masked by the same speaker. Same-talker segregation of speech is more challenging and effortful than different-talker segregation (<u>Brungart, 2001</u>) as it combines informational and energetic masking at spectral bands consistently defined by the same speaker’s vocal system. Therefore, under these conditions, priming may substantially assist the level of segregation needed for their particular task. Our one-trial design with varying speakers and voice combinations, by contrast, avoided the issue of speech mixtures’ learning effects, assessing listeners’ ability to segregate and understand continuous speech as in natural conditions. As a result, both studies have behaviorally distinct profiles where, in their case, target priming led to listener significant improvements in comprehension behavior relative to control conditions; in ours, such changes were not observed on already-proficient behavior.

The role that different levels of spectral clarity of the target’s degraded auditory signal play on listeners’ processing also matters from the perspective of target segregation. Varying spectral detail of an energetically masked speech stream produces substantially distinct response profiles in the auditory cortex, specifically the STS, under different levels of observers’ prior experience (Blank and Davis, 2015). Specifically, when listeners’ access to sensory detail in the speech stream is poor, providing clear prior expectations has the effect of increasing the quality of their STS multivariate representation of speech. If, on the other hand, speech input already has sufficient clarity, providing prior experience of similar clarity may by contrast lead to STS speech multivariate representations of reduced quality, in agreement with efficient coding principles. Hence, it would be possible that same-versus different-talker informational masking comparisons across the Wang et al. (2019) and the present study represent two levels of access to sensory detail about target speech that prior experience may differentially impact. Such opposing differences have been observed in the selectivity of multivariate neural responses (Blank and Davis, 2015) but possibly also in their univariate magnitude (cf. Turk-Browne et al., 2007 for visual scenes). The apparent discrepancy may then be summarized in the notion that weaker coding results from uninformative repetition, while coding enhancement follows the recognition of a stimulus achieved *through* repetition (Henson, 2003). Our reduced P2 findings appear furthermore consistent with findings of univariate response decrements by prior experience in single-speaker speech listening (Blank and Davis, 2015). Future studies may address specifically whether the spectrotemporal masking of speech similarly accounts for reversal effects of prior experience on selective neural representations in the cocktail party.

*Partial priming*. Speaker familiarity, defined by the long-term experience of a speaker’s voice, influences speech processing to the point of improving its perception. In certain cases, only short-term prior experience of unfamiliar voices improves behavior under masking (<u>Johnsrude et al., 2013</u>; <u>Daly and Pitt, 2023</u>). Semantic foreknowledge of speech content is a different form of behaviourally relevant prior experience in cocktail-party listening ( <u>Dekerle et al., 2014</u>). For instance, <u>Park et al.</u> (<u>2023</u>) studied the influence of topic familiarity on comprehension during attentional selection from speech mixtures. Familiar versus unfamiliar content was manipulated, along with presentations of fixed or variable speaker targets. When facing unfamiliar content (e.g., challenging philosophical discourse), listeners showed steep comprehension errors at rates that were not observed for familiar content, even under variable speaker voices. Such results are in support of the notion that sentential unpredictability impacts the level speech of intelligibility attainable under noisy conditions (<u>Bhandari et al., 2021</u>).

To evaluate whether the predictability of voice or content elements of speech impacts the coding of key envelope onset cues in multi-talker environments, we separately addressed these influences and examined them at each relevant processing stage. Voice and content cues relay spectral/prosodic and higher-level linguistic information, respectively, but may not convey contextual information specific to the mixture target’s temporal structure. Our present approach served to demonstrate the principle that when a given processing stage shows a reduced repetition effect to a target that only differs from a prime on, e.g., its temporal structure, then the processes subserved at that stage are sensitive to said structure (Henson, 2003). In other words, our partial priming manipulations of voice (Experiment 2a) or content (Experiment 2b) foreknowledge showed a lacking repetition effect on the P2 response to temporal edges. Failure to observe suppressed representations of voice-or content-primed targets here may be primarily interpreted in terms of listeners resolving target speech with an unpredictable envelope onset structure of the target signal. This demonstrates that despite its late latency, the P2 stage is also concerned with analysis of the temporal structure of the auditory target signal. Its absent repetition suppression when observer expectations no longer matched the target’s particular temporal structure indicates their role in the analysis of the unmasked target representation reached after the critical N1 stage. Responses at the P2 not only encode an unmasked neural estimate of speech but also include the listener’s expectations about when the voicing pattern of the target is being engaged.

## Conclusions

Multispeaker conditions present noisy and ambiguous challenges for the brain to define stable, invariant representations of incoming and relevant speech signals. Cortical tracking informs us about the brain’s ability to fixate on a single speaker nevertheless. How do neural synchronizations to key elements in the temporal structure of speech adjust as listener experience guides expectations? Top-down auditory attention triggers biased representations of these elements as early as 50 ms at the P1 stage, groups them as signal and noise components some 100 ms later at the N1, and defines its selection in some 100 ms more by the P2 stage. It is at this level that exact prior knowledge of selected speech modulates its auditory response by reducing it. Informing observers with only partial knowledge of speech, including the target speakers’ voice or speech contents, by contrast, does not lead to similar changes in envelope onset coding. Hence, P2 temporal edge coding at the cocktail party reflects the auditory brain’s undertaking of coordinating its activity to the tune of a neurally unmasked speech stream. The result, framed according to observer expectations gained through short-term learning, is an account of the temporal coordinates where to retrieve the most eventful acoustic information.

## Acknowledgments and funding sources

Conflict of interest: None to declare. We thank Franco Caballero, Rodrigo Caramés Harcevnicow, and Cecilia Biurrun for their assistance in data collection. We also thank Jonas Vanthornhout and Tom Francart for sharing their independent study data. The research presented in this manuscript received funding from the Agencia Nacional de Investigación e Innovación, Uruguay, under the scholarship POS_FCE_2020_1_1009198 to TSC and the research grant FCE_1_2019_1_155889 to FCC.

